# The N501Y spike substitution enhances SARS-CoV-2 transmission

**DOI:** 10.1101/2021.03.08.434499

**Authors:** Yang Liu, Jianying Liu, Kenneth S. Plante, Jessica A. Plante, Xuping Xie, Xianwen Zhang, Zhiqiang Ku, Zhiqiang An, Dionna Scharton, Craig Schindewolf, Vineet D. Menachery, Pei-Yong Shi, Scott C. Weaver

**Affiliations:** Department of Biochemistry and Molecular Biology, University of Texas Medical Branch, Galveston TX, USA; Institute for Human Infections and Immunity, University of Texas Medical Branch, Galveston TX, USA; World Reference Center for Emerging Viruses and Arboviruses, University of Texas Medical Branch, Galveston TX, USA; Department of Microbiology and Immunology, University of Texas Medical Branch, Galveston TX, USA; Texas Therapeutics Institute, Brown Foundation Institute of Molecular Medicine, The University of Texas Health Science Center at Houston, Houston, TX 77030, USA

## Abstract

Beginning in the summer of 2020, a variant of SARS-CoV-2, the cause of the COVID-19 pandemic, emerged in the United Kingdom (UK). This B.1.1.7 variant increased rapidly in prevalence among sequenced strains, attributed to an increase in infection and/or transmission efficiency. The UK variant has 19 nonsynonymous mutations across its viral genome including 8 substitutions or deletions in the spike protein, which interacts with cellular receptors to mediate infection and tropism. Here, using a reverse genetics approach, we show that, of the 8 individual spike protein substitutions, only N501Y exhibited consistent fitness gains for replication in the upper airway in the hamster model as well as primary human airway epithelial cells. The N501Y substitution recapitulated the phenotype of enhanced viral transmission seen with the combined 8 UK spike mutations, suggesting it is a major determinant responsible for increased transmission of this variant. Mechanistically, the N501Y substitution improved the affinity of the viral spike protein for cellular receptors. As suggested by its convergent evolution in Brazil and South Africa, our results indicate that N501Y substitution is a major adaptive spike mutation of major concern.

## Introduction

Since the emergence of severe acute respiratory syndrome coronavirus 2 (SARS-CoV-2) in 2019^1^, >100 million infections have occurred worldwide with >2.2 million fatalities. Despite some genetic proofreading during viral replication^2^, coronaviruses mutate at high frequencies^3^, leading to hundreds of mutations among SARS-CoV-2 strains. The trimeric spike protein consists of two subunits: the S1 subunit contains the receptor-binding domain (RBD) for angiotensin-converting enzyme 2 (ACE2), and the S2 subunit contains a peptide for membrane fusion^4^. Substitutions in the spike protein responsible for virus binding to ACE2 can strongly influence host range, tissue tropism, transmission, and pathogenesis^5^.

The first dominant mutation observed in SARS-CoV-2 encoded the spike protein D614G substitution^6^, which enhances viral replication in human airway epithelial cells and viral transmission in animal models^7–10^. Since then, additional spike mutations have been identified, including an N501Y spike substitution that occurred convergently in the United Kingdom (UK), South Africa, and Brazil. The UK variant B.1.1.7, which also contains 7 other spike substitutions, emerged in September, 2020. It was estimated to be 70-80% more transmissible than the ancestral lineage^11^ and has spread to more than 30 countries, including the United States^12,13^. Although B.1.1.7 variant strains and key mutants appear to be effectively neutralized by vaccine-induced antibodies^14–16^, their increased transmissibility could counteract increasing herd immunity due to vaccination and natural infections. Thus, B.1.1.7 threatens to exacerbate the COVID-19 pandemic as it continues to spread and displace earlier variants.

To investigate potential mechanisms of increased transmission of the B.1.1.7 variant, we used reverse genetics, the hamster model, and primary human airway epithelial cell cultures to probe the impact of the spike mutations found in B.1.1.7. We determined the phenotypes of individual spike mutations, as well as that of the 8 combined mutations (UK-8x), using competition fitness assays to compare mutants to the ancestral G614 strain. Our results suggest that, consistent with its convergent evolution, the N501Y substitution is a critical spike protein determinant of enhanced infection of the upper airway and transmission.

## Results

To investigate phenotypes of the 8 spike amino acid changes found in the UK B.1.1.7 variant, we infected golden Syrian hamsters intranasally with SARS-CoV-2 mutants encoding each of the individual spike substitutions, as well as the combination of all 8 (UK-8x; Extended Data Fig. 1a,b). These mutations were engineered into the USA-WA1/2020 strain^17^ containing the D614G spike substitution that has become dominant worldwide due to its increased transmission efficiency^8^; the resulting SARS-CoV-2 strain [USA-WA1/2020-G614, hereafter called wild-type (wt)], was generated using site-directed mutagenesis of a cDNA clone^18^. There was no major effect of the mutations on plaque phenotypes on Vero cells (Extended Data Fig. 1c). Each individual mutant and the combination UK-8x were tested in an *in vivo* competition assay by mixing an approximately 1:1 ratio of mutant:wt prior to intranasal infection of hamsters (Fig. 1a). This competition assay has major advantages compared to individual strain infections: the replication of each virus in the competition is internally controlled, eliminating host-to-host variation that can reduce experimental power, and the ratios of the virus strains can be assayed with more precision than individual virus titers. Thus, competition assays have been used for many studies of microbial fitness^19–22^ including SARS-CoV-2^8^, due to their precision and reproducibility^23^.

**Figure 1.**
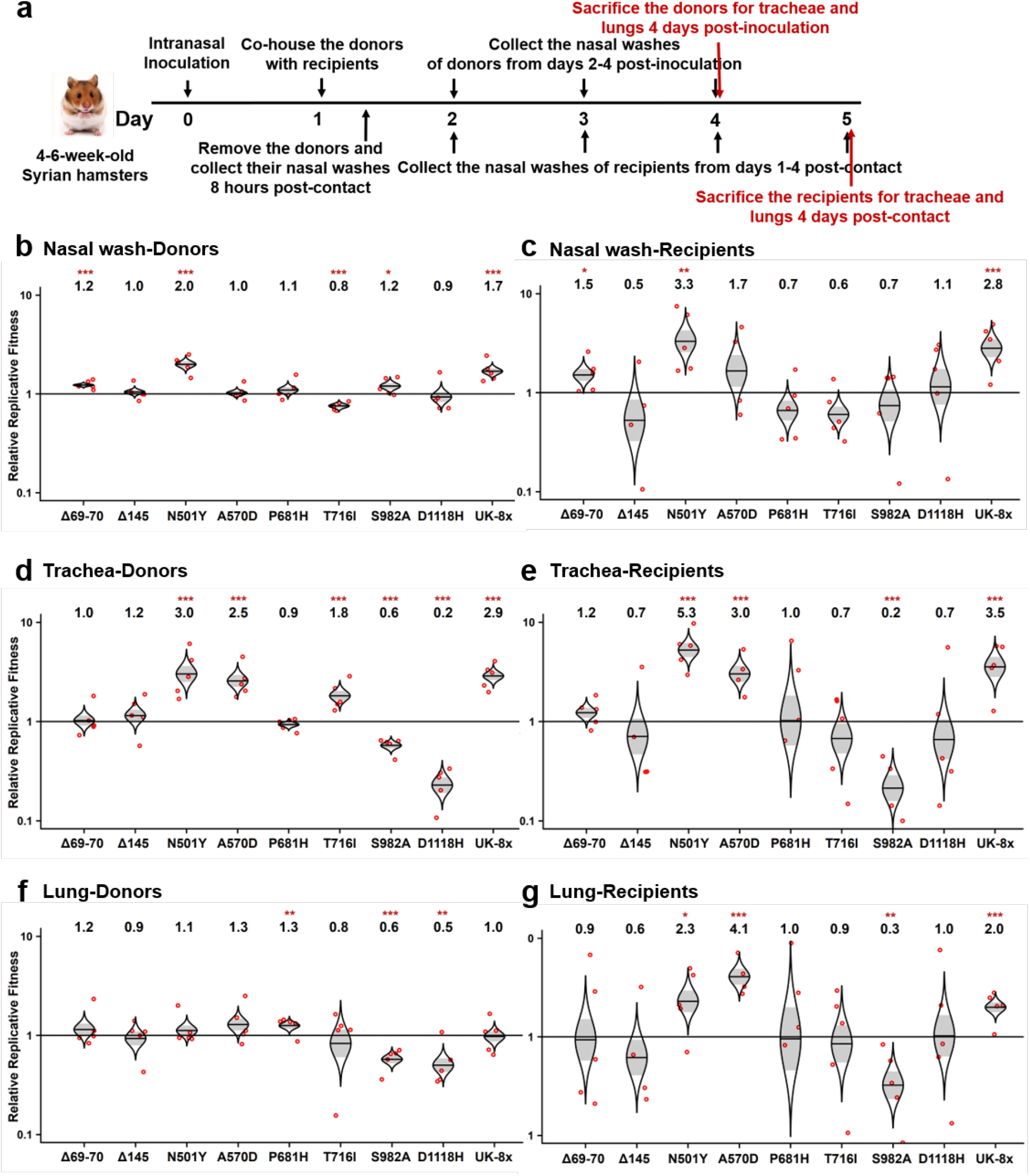
The screening of the SARS-CoV-2 UK variant spike substitutions in hamsters by competition assay. **a,** Design of the hamster competition fitness studies. The mutant viruses were mixed with wild type G614 virus and inoculated into donor hamsters intranasally (I.N.) at a total titer of 10^5^ PFU per hamster. The donor hamsters were co-housed with recipient hamsters 1 day post-infection. After 8 hours of contact, the donors were removed. All hamsters were subjected to nasal washes daily 1-4 days after infection and sacrificed for organ collections 4-days post-inoculation or post-contact. **b-g,** The competition of different UK variants with wild-type (wt) G614 virus. **b,d,f,** Final:inoculum ratios of 8 individual UK variants and the UK-8x variant (containing all 8 spike mutations) with the wt in nasal washes (**b**), tracheae (**d**) and lungs (**f**) of donor hamsters 4 days post-inoculation. **c,e,g,** Final:donor inoculum ratios of 8 individual UK mutations and UK-8x variant with the wt in nasal washes (**c**), tracheae (**e**) and lungs (**g**) of recipient hamsters 4-days post-contact. **b-g,** Red dots represent individual animals (n = 5), the horizontal lines in each catseye represent the mean, shaded regions represent standard error of the mean; y-axes use a log10 scale. Black numbers above each set of values (catseye) indicate the relative fitness estimates. P values are calculated for the group (strain) coefficient for each linear regression model. *p<0.05; **p<0.01; ***p<0.001.

Hamsters were first sampled 4 days post-infection after the peak of viral loads^8^, using nasal washes and necropsied tracheae and lungs (Fig. 1a). One day after infection, the animals were co-housed with recipient hamsters for 8 hr of direct contact to assess transmission. The recipients were held an additional 4 days for equivalent sampling. For the inocula and hamster samples, RT-PCR reactions with primers spanning the mutations were performed, and the amplicon DNAs underwent Sanger sequencing and electropherogram peak analyses to determine mutant:wt ratios. Ratio changes in the hamster samples compared to the inoculum reflect of the relative fitness advantage for the winner of the competition.

The competition results assessed from nasal washes indicated that, of the 8 mutations examined, only the deletion of codons 69-70 (Δ69-70), N501Y, S982A, and the combined UK-8x conferred significant fitness advantages, with other substitutions showing slightly but inconsistently reduced fitness or no difference from wt (Fig. 1b). In recipient samples, the same overall fitness trends were observed (Fig. 1c). Examination of the tracheal samples showed similar results (Fig. 1d), except that Δ69-70 no longer impacted fitness, while A570D increased and two other mutations (S982A and D1118H) decreased fitness. Following transmission, the same overall fitness effects were maintained in the tracheae (Fig. 2e), consistent with findings from nasal washes (Fig. 1c). Notably, results from lung samples collected at day 4 were distinct from those in the upper airway (nasal washes and tracheae), with only slight fitness effects of the spike mutations; even the combination UK-8x did not increase fitness significantly for donor replication in the lungs (Fig. 1f). Yet, significant fitness gains were observed in recipients for N501Y and A570D as well as UK-8x, suggesting more efficient transmission of these variants, and significant losses were maintained only for S982A (Fig. 1g). Overall, these experiments demonstrated that only the N501Y substitution conferred a consistent, significant, major fitness benefit in the upper airway, comparable to UK-8x.

In the subsequent study, we found that N501Y also conferred an advantage across both early and late times of nasal shedding. Evaluating nasal washes on days 1-4, N501Y conferred a significant fitness advantage beginning on day 3 in the donors, and this advantage was maintained in day 1-4 recipient washes (Fig. 2a,b). Evaluation of the other substitutions in nasal washes collected 1-4 days after infection or following transmission demonstrated that only Δ69-70 and UK-8x showed consistent fitness gains (Extended Data Fig. 2). Total viral loads in hamster samples were similar in donor and recipient animals (Extended Data Fig. 3), although nasal washes were consistently lower in recipients on day 1, probably reflecting a lower infection dose reaching recipients from intra-cage transmission than from direct intranasal inoculation via donors. Altogether, these results demonstrate a clear fitness advantage conferred by N501Y for shedding in the upper airway, including after transmission.

**Figure 2.**
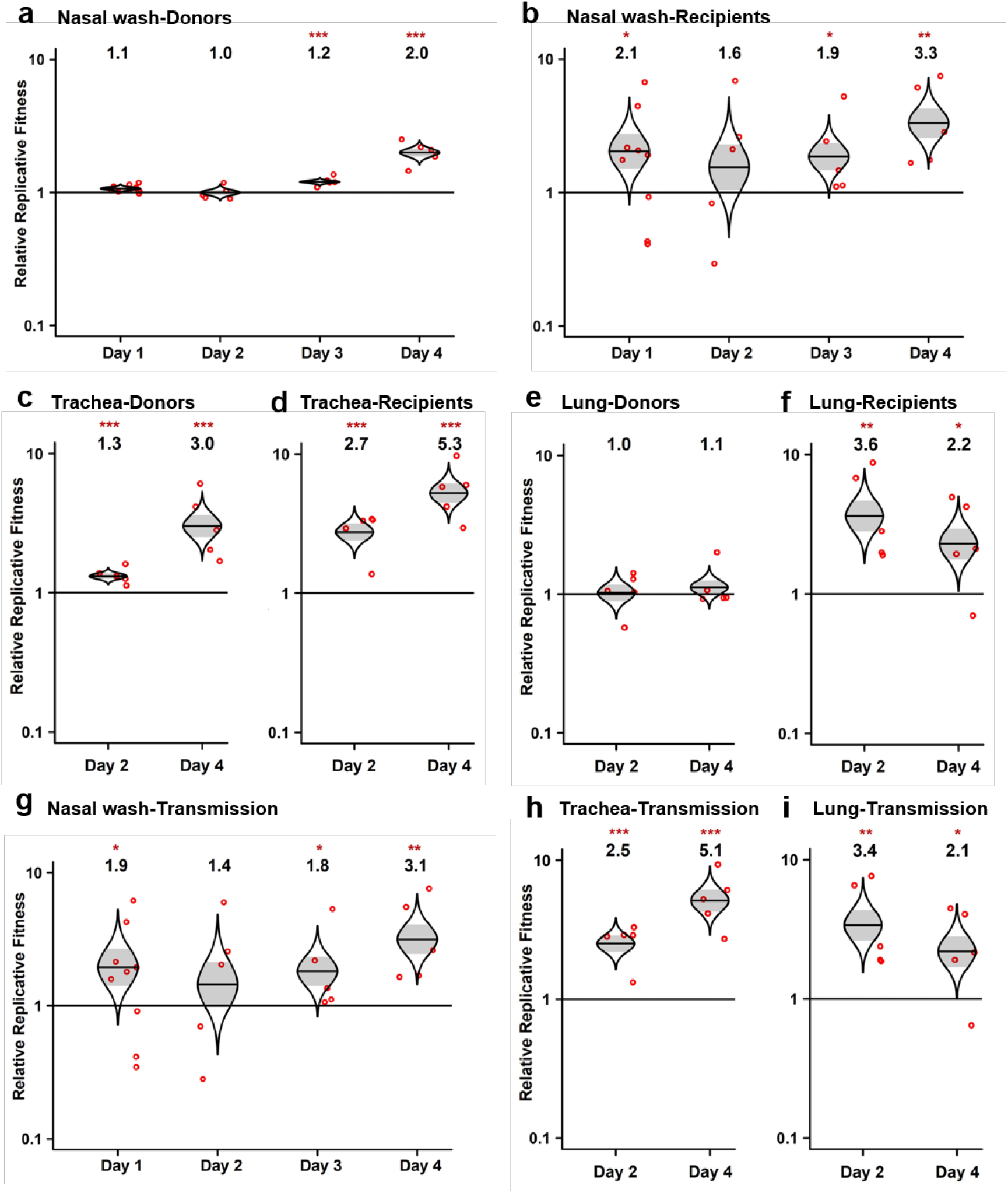
The SARS-CoV-2 spike N501Y mutant has a consistent advantage in competitions with wt during the upper airway replication and transmission between hamsters. **a,b,** Results of the competition between spike N501Y mutant and the wt assessed by sampling nasal washes of both donor (**a**) and recipient hamsters (**b**) from 1-4 days post-inoculation (donors) or post-contact (recipients). **c,d,** Results of the competition between the N501Y mutant and the wt in the tracheae of both donor (**c**) and recipient hamsters (**d**) at 2- or 4-days post-inoculation or post-contact. **e,f,** Results of the competition between the N501Y mutant and the wt in the lungs of both donor (**e**) and recipient hamsters (**f**) sampled 2- or 4-days post-inoculation or post-contact. **g-i,** The ratios in the competitors in the nasal washes (**g**), tracheae (**h**) and lungs (**i**) of recipient hamsters were compared to the ratio of N501Y:wt measured from the day 1 nasal wash of donor hamsters (representing the virus population transmitted) to assess changes corresponding to transmission versus replication in the recipient hamsters. **a-i,** The fitness advantage of the N501Y substitution against wt both during infection (donor data) and after transmission of the virus to recipients is shown by the significant changes in ratios between the harvested samples and inocula. Red dots represent individual animals (n=5), the horizontal lines in each catseye represent the mean, shaded regions represent standard error of the mean; y-axes use a log_10_ scale. Black numbers above each set of values (catseye) indicate the relative fitness estimates. P values are calculated for the group (strain) coefficient for each linear regression model. *p<0.05; **p<0.01; ***p<0.001.

Because it conferred the major phenotype among the 8 spike gene mutations, we next examined N501Y in tracheal necropsies sampled on days 2 and 4 days in both donors and recipients. Similar to nasal washes, the tracheal samples showed a consistent fitness advantage of N501Y at both timepoints (Fig. 2c,d). Calculations of fitness in the recipient hamsters after transmission were also performed using the donor day 1 nasal wash ratio, representing the population transmitted, as the starting ratio rather than the original donor inoculum; this had little effect on fitness estimates (Fig. 2g-i). The UK-8x mutant also showed a consistent, significant fitness advantage over the wt after transmission in both nasal washes (Extended Data Fig. 4a) as well as tracheae and lungs harvested on day 4 after infection (Extended Data Fig. 4b). Overall, our in vivo experiments revealed a consistent fitness advantage of N501Y and UK-8x, and to a lesser extent Δ69-70, for replication in the upper hamster airway.

To examine the fitness effect of the UK spike substitutions on human and other primate cell lines, we performed both replication kinetics with individual SARS-CoV-2 mutants, and competition assays using Vero E6 and Calu-3 cells. In Vero cells sampled 12-48 hr post-infection, UK-8x and N501Y consistently replicated to higher titers than the wt as measured using infectious plaque assays (Extended Data Fig. 5a). There were no major differences among the 3 strains in Calu-3 cells, although N501Y and UK-8x had a trend of faster replication that was not statistically significant (Extended Data Fig. 5b). Measurement of viral titers with RT-qPCR gave similar results (Extended Data Fig. 5c,d). Only in Vero cells were there consistent differences in genomic RNA:plaque-forming unit (PFU) ratios, with the wt higher at several time points (Extended Data Fig. 5e,f).

In primary human airway epithelial (HAE) cultures (Fig. 3a), the N501Y and UK-8x mutants replicated significantly faster in early stages of infection, as measured in PFU (Fig. 3c), but there was little difference when measured in RNA copies (Fig. 3d). The RNA:PFU ratios were significantly lower for N501Y and UK-8x compared to wt on days 1-3, suggesting greater specific infectivity of these mutants. In mixed competition infections, N501Y showed significantly higher fitness at nearly all time points in Vero and Calu-3 cells (Fig. 3e,f). In HAE cells, both N501Y (Fig. 3g) and UK-8x (Extended Data Fig. 6) showed major fitness advantages over wt (Fig. 3g), consistent with our *in vivo* data, suggesting that this substitution is a major determinant of the fitness advantage of the UK variant over wt SARS-CoV-2.

**Figure 3.**
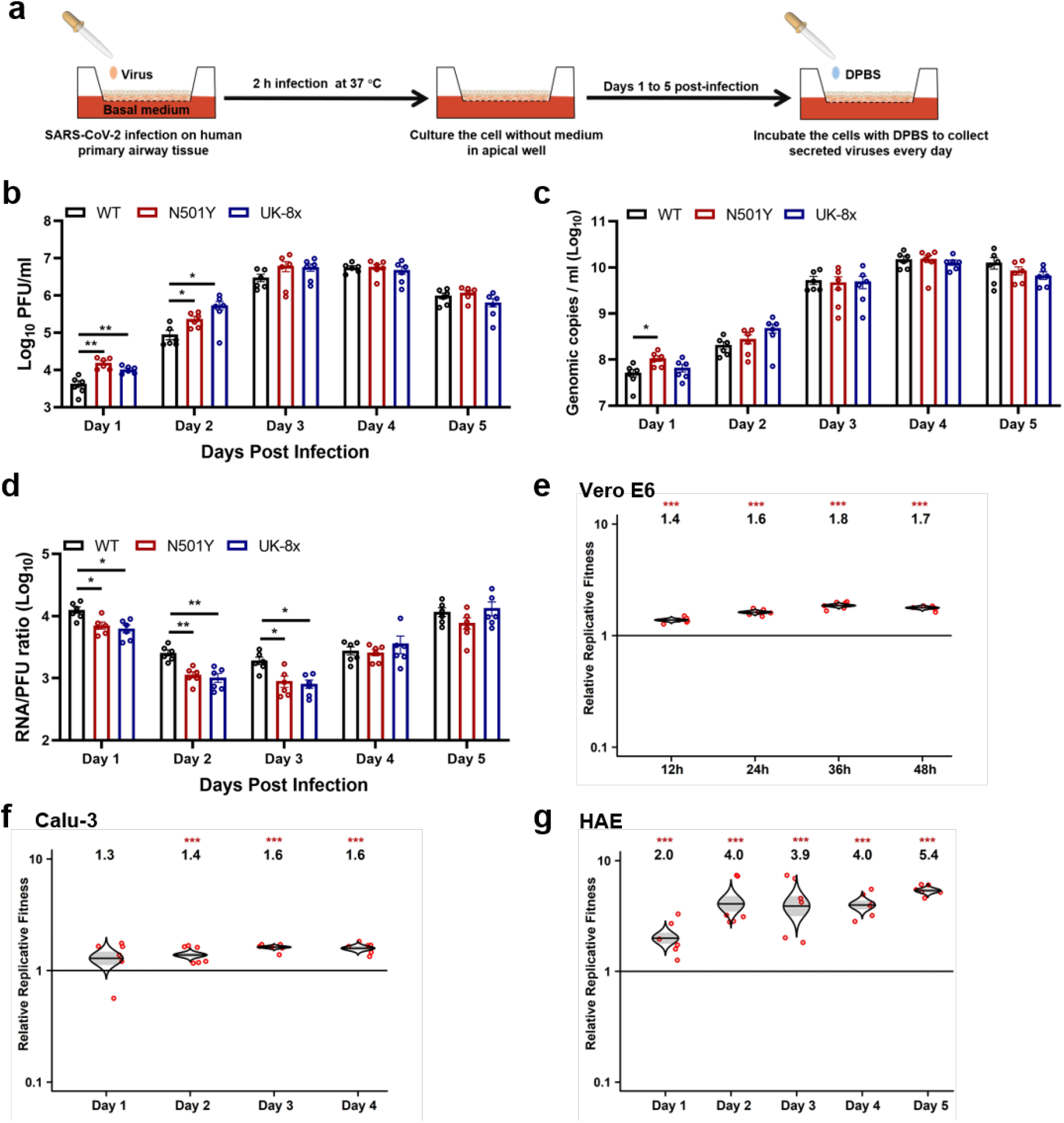
The spike N501Y mutation enhances viral replication in primary human airway cells and has advantages in the competition against wt virus *in vitro*. **a,** The experimental scheme of primary human airway cell infections. The wt, N501Y and UK-8x mutants were inoculated onto human airway epithelial (HAE) cells at a MOI of 5. After a 2 h incubation, the culture was washed with DPBS and maintained for 5 days. The secreted viruses were collected in DPBS after 30 min incubation at 37°C every day. **b-d,** The replication kinetics of the N501Y and UK-8x mutants compared to the wt on HAE cells. The amounts of infectious virus (**b**) and genomic RNA (**c**) were quantified by plaque assay and RT-qPCR, respectively. The genomic RNA:PFU ratio (**d**) was calculated as an indication of virion infectivity. The detection limitation of the plaque assay was 10 PFU/ml. Dots represent individual biological replicates (n=6) pooled from two independent experiments. The values in the graph represent the mean ± standard error of the mean. A non-parametric Mann-Whitney test was used to determine significant differences. P values were adjusted using the Bonferroni correction to account for multiple comparisons. Differences were considered significant if *p<0.025; **p<0.005. **e-g,** The Competition assay between N501Y variant and wt virus on Vero E6 (**e**), calu-3 (**f**) and HAE (**g**) cells. The Vero E6, calu-3 and HAE cells were infected at a MOI of 0.01, 0.1 and 5, respectively. Red dots represent individual animals (n=5), the horizontal lines in each catseye represent the mean, shaded regions represent standard error of the mean; y-axes use a log10 scale. Black numbers above each set of values (catseye) indicate the relative fitness estimates. P values are calculated for the group (strain) coefficient for each linear regression model. ***p<0.001.

To assess potential effects on virulence, individual mutant infections of hamsters were examined for weight loss, with no significant differences observed between wt, N501Y and the UK-8x mutant (Extended Data Fig. 7a,b). This suggests that the UK spike mutations may not affect virulence. Nasal washes of these infected animals showed consistently higher titers on days 1-3 and 5 for both N501Y and UK-8x, but only when measured in PFU. However, these differences were significant on only on day 1 (Extended Data Fig. 7c,d), again reflecting the lesser sensitivity of individual infections compared to competition assays to detect small fitness differences. Genomic RNA:PFU ratios were consistently higher for the wt than the mutants (Extended Data Fig. 7e), again suggesting greater specific infectivity of the latter. Trachea and lung samples showed no significant titer differences when measured as PFU or RNA copies (Extended Data Fig. 7f,g), and the RNA:PFU ratio was higher for wt on day 1 (Extended Data Fig. 7h). Together, these results again suggest a fitness benefit for N501Y replication and transmission from the upper airways in the hamster model of COVID-19.

As the SARS-CoV-2 pandemic has progressed, several mutations have been convergently selected in variant lineages; of the 8 spike substitutions seen in the UK variant, only Δ69-70 and N501Y have evolved convergently in other variants, consistent with our fitness results (Fig. 4a). The frequency of the N501Y mutation worldwide has increased dramatically since October, 2020 (Fig. 4b). Coupled with our *in vivo* and *in vitro* competition results, these data indicate that the N501Y substitution provides a major fitness advantage to variants that maintain it.

**Figure 4.**
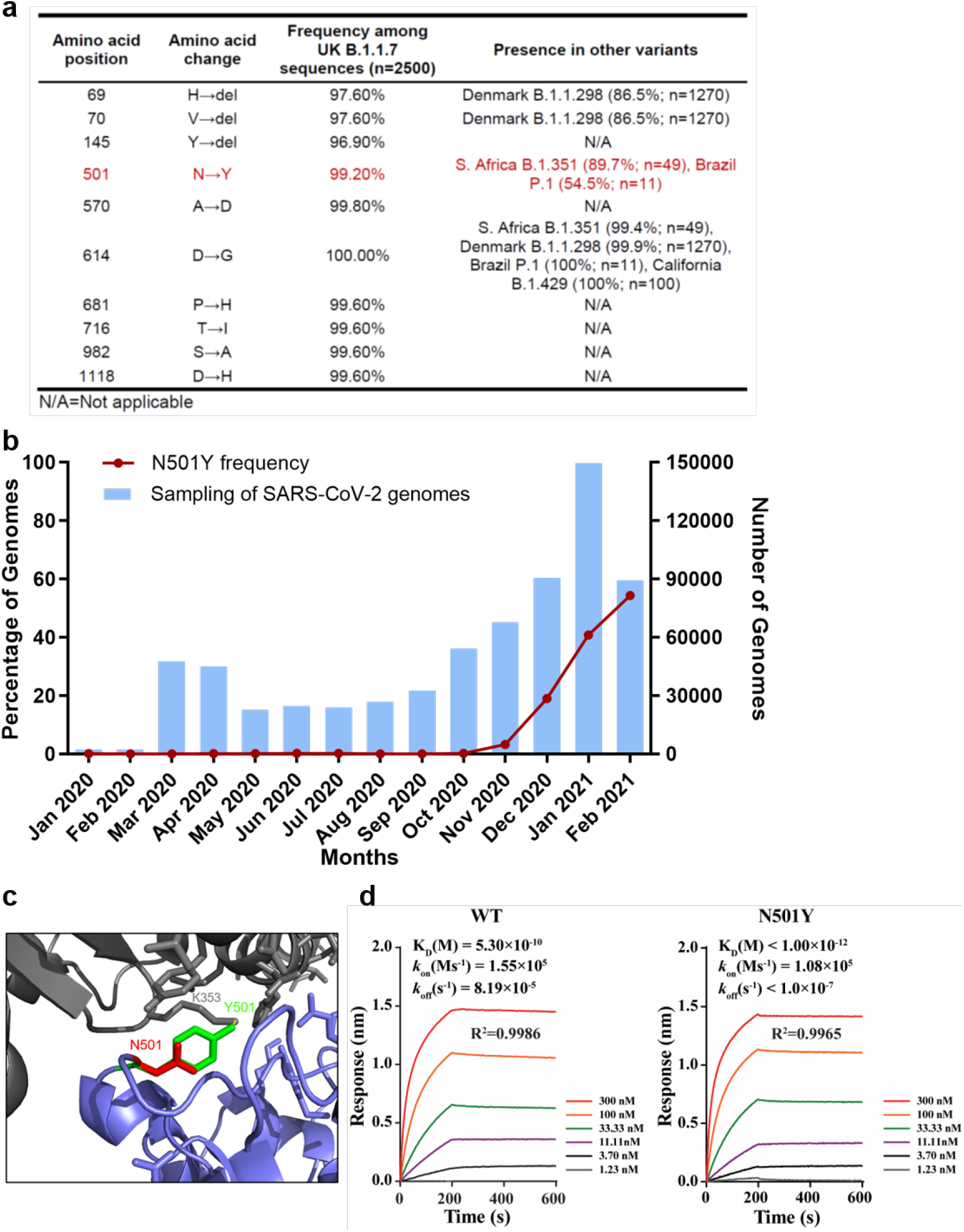
The spike N501Y substitution spread quickly and increases spike protein binding affinity for the human ACE2 receptor. **a,** The frequencies of the spike protein amino acid substitutions found among UK B.1.1.7 isolates and other variants. The amino acid substitutions in the UK B.1.1.7 variant refers to the USA_WA1/2020 SARS-CoV-2 sequence (GenBank accession No. MT020880). **b,** The frequency of the N501Y substitution over time in all genomic SARS-CoV-2 sequences available from the GISAID database worldwide. The blue bars represent the total numbers of SARS-CoV-2 genomes sequenced worldwide. The red line indicates the percentage of N501Y variant in total SARS-CoV-2 genomes. **c,** The predicted binding site of the N501 and Y501 residues to the human ACE2 receptor. **d,** The binding affinities of the wt and N501Y mutant to the human ACE2 receptor. Binding affinity-related parameters, including association (K_on_), dissociation (K_off_), and affinity (K_D_) are shown. The affinity of ACE2 to the N501Y mutant RBD is below the detection limit and is presented as <10^−12^.

We next examined the mechanism for this fitness advantage, focusing on the impact of N501Y on receptor binding. Located in the spike RBD (Fig. 4c), the change of asparagine to tyrosine is expected to increase interactions with the human ACE2 receptor at residue 353, leading to increased binding affinity. To address this hypothesis, we conducted binding assays using recombinant spike RBD and human ACE2 proteins on a Bio-Layer Interferometry (BLI) system (Fig. 4d). The N501Y substitution increased the RBD/ACE2 binding, as indicated by the >350-fold and >819-fold improved K_D_ and K_off_, respectively. These results suggest that the N501Y substitution improves viral fitness for replication in the upper airway, resulting in enhanced transmission, via an enhanced spike/receptor interaction.

## Discussion

To corroborate epidemiologic findings that the UK B.1.1.7 SARS-CoV-2 variant is more efficiently transmitted^11^, and to identify genetic determinants of this phenotype, we performed experimental infections of hamsters, cell lines, and primary human airway epithelial cells with each of the 8 individual spike gene mutations. Only the N501Y substitution and a deletion of codons 69-70 showed consistent fitness advantages for replication in the upper airway in the hamster model, with higher shedding in nasal secretions, as well as in primary human airway epithelial cells. The N501Y substitution alone had a phenotype similar to that of the combined 8 mutations, suggesting it is the major spike determinant driving increased transmission of the UK variant. This substitution, also present in the South African and Brazilian variants, is likely a result of convergent evolution, suggesting that N501Y is a major adaptive spike mutation of current concern. The mechanism of enhanced fitness is likely related to increased replication and shedding in the nasal cavity, leading to more efficient air-borne and/or fomite-mediated transmission. Fortunately, tests to date with N501Y substitution indicate minimal changes in susceptibility to in PRNT_50_ values^14–16^ suggesting little or no resistance to neutralization elicited by vaccines now in widespread use. These along with our data suggest UK variant adaptation for more efficient receptor binding rather than antibody escape..

Although Δ69-70 has also been found in a Denmark mink variant (Fig. 4a) as well as in wt SARS-CoV-2 strains in several locations, it has not evolved convergently in the major variants with putative enhanced transmission. Furthermore, in the United States, despite its presence since October, 2020, it has not increased dramatically in frequency (Larsen and Worobey, https://virological.org/t/identification-of-a-novel-sars-cov-2-spike-69-70-deletion-lineage-circulating-in-the-united-states/577). This finding corroborates our results indicating that this mutation has less impact on transmission potential than N501Y, as indicated by our *in vivo* and *in vitro* models. However, the frequent co-occurrence of this deletion^24^ with N501Y suggests possible epistasis, which should be further examined.

Considering that only 2 of the 8 spike substitutions conferred fitness advantages for infection of the upper airway and transmission, the origins and roles of the other 6 deserve consideration. The first of these to appear in sequenced strains were the two deletions (69-70 and 145), followed by all of the substitutions except S982A by March, 2020 (Extended Data Table 1). However, beginning in September, 2020, all 8 mutations, but especially N501Y (Extended Table 1 and Fig. 4b), began increasing dramatically in prevalence and almost simultaneously in frequency. Combined with our data indicating that none except Δ69-70 and N501Y has a consistent phenotype, and some even show reduced fitness in our experimental results at certain timepoints, this suggests many if not all of the remaining mutations were not directly selected; these mutations probably occurred through drift mechanisms such as founder and hitchhiking effects, followed by maintenance via linkage to Δ69-70 and N501Y, or possibly via recombination, which occurs frequently during SARS-CoV-2 replication^3^. Other possibilities include epistatic interactions among these mutations that were not revealed via our analysis of individual mutations or the combination of all 8. It is also possible that mutations in other parts of the genome contribute to the overall fitness of the UK variant for replication and transmission.

Another limitation of our study is the models used. The hamster may be the best rodent model available^8^, but its ACE2 receptors differ slightly from those of humans^25^. Although we combined the hamster model with infection of primary human airway cells to address this limitation, the use of a primate model would be ideal to confirm our findings.

In conclusion, we have used authentic SARS-CoV-2 to demonstrate that only 2 of 8 spike substitutions in the UK B.1.1.7 variant are mainly responsible for the enhanced transmission of this strain. Although mutations in other parts of the genome could also contribute to this transmission phenotype, the N501Y substitution in the spike protein in particular appears to be a major determinant of efficient transmission. This mutation is present in several regions of the world and should be closely monitored to anticipate public health measures needed to control SARS-CoV-2 spread.

## Methods

### Ethics statement

Hamster studies were performed in accordance with the guidance for the Care and Use of Laboratory Animals of the University of Texas Medical Branch (UTMB). The protocol was approved by the Institutional Animal Care and Use Committee (IACUC) at UTMB. All the hamster operations were performed under anesthesia by isoflurane to minimize animal suffering.

### Animals and Cells

The Syrian golden hamsters (HsdHan:AURA strain) were purchased from Envigo (Indianapolis, IN). African green monkey kidney epithelial Vero E6 cells (provided by R. Baric) were grown in Dulbecco’s modified Eagle’s medium (DMEM; Gibco/ThermoFisher, Waltham, MA, USA) with 10% fetal bovine serum (FBS; HyClone Laboratories, South Logan, UT) and 1% antibiotic/streptomycin (Gibco). Human lung adenocarcinoma epithelial Calu-3 cells (ATCC, Manassas, VA, USA) were maintained in a high-glucose DMEM containing sodium pyruvate and GlutaMAX (Gibco) with 10% FBS and 1% penicillin/streptomycin at 37°C with 5% CO_2_. The EpiAirway system is a primary human airway 3D tissue model purchased from MatTek Life Science. This EpiAirway system was maintained with the provided culture medium at 37 °C with 5% CO_2_ following the manufacturer’s instruction. All other culture medium and supplements were purchased from ThermoFisher Scientific (Waltham, MA). All cell lines were verified and tested negative for mycoplasma.

### Generation of SARS-CoV-2 mutant viruses

Individual point mutations in the spike gene (Δ69-70, Δ145, N501Y, A570D, P681H, T716I, S982A, D1118H and UK-8x) were introduced into an D614G infectious cDNA clone of the 2019n-CoV/USA_WA1/2020 (WA1/2020) strain as described previously^26^. Briefly, the nucleotide substitutions were introduced into a subclone puc57-CoV-2-F5-7 containing the spike gene of the SARS CoV-2 wild-type infectious clone^18^ by overlap fusion PCR. The full-length infectious cDNA clones of the variant SARS-CoV-2 viruses were assembled by *in vitro* ligation of contiguous cDNA fragments following the protocol previously described^18^. *In vitro* transcription was then preformed to synthesize full-length genomic RNA. For recovering the mutant viruses, the RNA transcripts were electroporated into Vero E6 cells. The viruses from electroporated cells were harvested at 40 h post-electroporation and served as p0 stocks. All viruses were passaged once on Vero E6 cells for subsequent experiments and subjected to sanger sequencing after RNA extraction to confirm the introduction and stability of substitutions (Extended Data Table 2).Viral titers were determined by plaque assay on Vero E6 cells. All virus preparation and experiments were performed in a biosafety level 3 (BSL-3) facilities. Viruses and plasmids are available from the World Reference Center for Emerging Viruses and Arboviruses at the University of Texas Medical Branch.

### RNA extraction, RT-PCR, and Sanger sequencing

Cell culture supernatants or clarified tissue homogenates were mixed with a five-fold excess of TRIzol™ LS Reagent (Thermo Fisher Scientific, Waltham, MA). Viral RNAs were extracted according to the manufacturer’s instructions. The extracted RNAs were dissolved in 20 μl nuclease-free water. Two microliters of RNA samples were used for reverse transcription by using the SuperScript™ IV First-Strand Synthesis System (ThermoFisher Scientific) with random hexamer primers. Nine DNA fragments flanking the entire viral genome were amplified by PCR. The resulting DNAs were cleaned up by the QIAquick PCR Purification Kit, and the genome sequences were determined by Sanger sequencing at GENEWIZ (South Plainfield, NJ).

The quantify viral RNA samples, quantitative real-time RT-PCR assays were performed using the iTaq SYBR Green One-Step Kit (Bio-Rad) on the LightCycler 480 system (Roche, Indianapolis, IN) following the manufacturers ’protocols. Primers are listed in Extended Data Table 2. The absolute quantification of viral RNA was determined by a standard curve method using an RNA standard (*in vitro* transcribed 3,480 bp containing genomic nucleotide positions 26,044 to 29,883 of SARS-CoV-2 genome).

To quantify mutant ratios for competition assays, RT-PCR products were amplified from extracted RNA using a SuperScript™ III One-Step RT-PCR kit (Invitrogen, Carlsbad, CA, USA). A 20 μl reaction was assembled in PCR 8-tube strips through the addition of 10 μl 2× reaction mix, 0.4 μlSuperScript III RT/Platinum Taq Mix, 0.8 μl Forward Primer (10 μM) (Extended Data Table 2), 0.8 μl reverse primer (10 μM), 4 μl RNA, and 6 μl Rnase-free water. Reverse transcription and amplification was completed using the following protocol: (i) 55°C, 30 min; 94°C, 2 min; (ii) 94°C, 15s; 60°C, 30s; 68°C, 1min; 40 cycles; (iii) 68°C, 5 min; (iv) indefinite hold at 4°C. The presence and size of the desired amplicon was verified with 2 μl of PCR product on an agarose gel. The remaining 18 μl were purified by a QIAquick PCR Purification kit (Qiagen, Germantown, MD) according to the manufacturer’s protocol.

Sequences of the purified RT-PCR products were generated using a BigDye Terminator v3.1 cycle sequencing kit (Applied Biosystems, Austin, TX, USA). The sequencing reactions were purified using a 96-well plate format (EdgeBio, San Jose, CA, USA) and analyzed on a 3500 Genetic Analyzer (Applied Biosystems, Foster City, CA).The peak electropherogram height representing each mutation site and the proportion of each competitor was analyzed using the QSVanalyser program^27^.

### Plaque assay

Approximately 1.2×10^6^ Vero E6 cells were seeded to each well of 6-well plates and cultured at 37°C, 5% CO_2_ for 16 h. Virus was serially diluted in DMEM with 2% FBS and 200 μl diluted viruses were transferred onto the monolayers. The viruses were incubated with the cells at 37°C with 5% CO_2_ for 1 h. After the incubation, overlay medium was added to the infected cells per well. The overlay medium contained DMEM with 2% FBS, 1% penicillin/streptomycin and 1% sea-plaque agarose (Lonza, Walkersville, MD). After a 2-days incubation, plates were stained with neutral red (Sigma-Aldrich, St. Louis, MO) and plaques were counted on a light box.

### Viral infection of cell lines

Approximately 3×10^5^ Vero E6 or calu-3 cells were seeded onto each well of 12-well plates and cultured at 37°C, 5% CO_2_ for 16 h. The infection titers for Vero E6 and calu-3 cells were 0.01 and 0.1 MOI, respectively. The virus was incubated with the cells at 37°C for 2 h. After infection, the cells were washed with DPBS 3 times to remove any unattached virus. One milliliter of culture medium was added into each well for the maintenance of the cells. At each time point, 150 μl of culture supernatants were harvested for the real-time qPCR detection and plaque assay, and 150 μl fresh medium was added into each well to replenish the culture volume. The cells were infected in triplicate for each virus. All samples were stored at −80°C until plaque or RT-PCR analysis.

### Viral infection in a primary human airway cell culture model

The EpiAirway system is a primary human airway 3D mucociliary tissue model consisting of normal, human-derived tracheal/bronchial epithelial cells. For viral replication kinetics, WT, N501Y or UK-8x mutant viruses were inoculated onto the culture at a MOI of 5 in DPBS respectively. After 2 h infection at 37 °C with 5% CO_2_, the inoculum was removed, and the culture was washed three times with DPBS. The infected epithelial cells were maintained without any medium in the apical well, and medium was provided to the culture through the basal well. The infected cells were incubated at 37 °C, 5% CO_2_. From 1-5 days, 300 μl of DPBS were added onto the apical side of the airway culture and incubated at 37 °C for 30 min to elute the released viruses. All virus samples in DPBS were stored at −80°C.

### Spike RBD and ACE2 binding

The human ACE2 protein was purchased from Sino Biological (Beijing, China; Cat# 10108-H08H) and the human IgG1 Fc-tagged RBD proteins were made in-house using a method as previously described^28^. The affinity measurement was performed on the Forte bio Octet RED 96 system (Sartorius, Goettingen, Germany). Briefly, wt or N501Y mutant RBD proteins (20μg/ml) were captured onto protein A biosensors for 300s. The loaded biosensors were then dipped into the kinetics buffer for 10s for adjustment of baselines. Subsequently, the biosensors were dipped into serially diluted (1.23~300nM) human ACE2 protein for 200s to record association kinetics and then dipped into kinetics buffer for 400s to record dissociation kinetics. Kinetic buffer without ACE2 was used to correct the background. The Octet Data Acquisition 9.0 software was used to collect affinity data. For fitting of K_D_ values, Octet Data Analysis software V11.1 was used to fit the curve by a 1:1 binding model and use of the global fitting method.

### Hamster infections

Four- to six-week-old male golden Syrian hamsters, strain HsdHan:AURA (Envigo, Indianapolis, IN), were inoculated intranasally with 100 μl SARS-CoV-2. For transmission competition assays, five donor animals received a mixture containing 5 x 10^4^ PFU of G614 virus (wt) and an equal amount of variant virus. One day later, one infected donor animal was co-housed with one naïve animal for 8 hours (10 pairs for N501Y group, 5 pairs for the other groups) and the donors were returned to their cages. For virus replication assays, the animals received DMEM with 2% FBS and 1% penicillin/streptomycin (Mock, n=4), wt G614 virus (n=9), N501Y mutant virus (n=9), UK-8x mutant virus (n=9) at a dose of 10^4^ PFU/hamster. The infected animals were weighed and monitored for signs of illness daily. Nasal washes were collected in 400 μl sterile DPBS at indicated time points. Animals were humanely euthanized for organ collections at 2- or 4 days post-inoculation (donors) or post-contact (recipients) for the transmission competition study. In the virus replication study, 4 hamsters in each group were sacrificed for organ collection 2 days post-infection and others were euthanized 7 days post-infection. The harvested tracheae and lungs were placed in a 2-ml homogenizer tube containing 1 ml of maintenance media (DMEM supplemented with 2% FBS and 1% penicillin/streptomycin) and stored at −80°C. Samples were subsequently thawed, lung or tracheae were homogenized for 1 min at 26 sec-1, and debris was pelleted by centrifugation for 5 min at 16,100×g. Infectious titers were determined by plaque assay. Genomic RNAs were quantified by quantitative RT-PCR (primers in Extended Data Table 2). Ratios of mutant:wt RNA were determined via RT-PCR with quantification of Sanger peak heights.

### Competition assays

For the competition assay on Vero E6, calu-3 and primary human epithelial airway cells, WT and mutant viruses were mixed and inoculated onto the cells at a final MOI of 0.01, 0.01 and 5 respectively. For the competition in hamsters, 100 μl mixtures of wt and variant viruses (total 1×10^5^ PFU per hamster) were inoculated intranasally into 4-6-week-old Syrian hamsters. On 1-4 days post-inoculation (donors) or post-contact (recipients), the infected hamsters were sampled for competition detection. An aliquot of the inoculum used for both hamster and cell infections was back-titered to confirm the initial ratio of viruses. All samples were stored in −80°C freezer prior to analysis.

### Statistics

Male hamsters were randomly allocated into different groups. The investigators were not blinded to allocation during the experiments or to the outcome assessment. No statistical methods were used to predetermine sample size. Dead animals were excluded from sample collections and data analysis. Descriptive statistics have been provided in the figure legends. For *in vitro* replication kinetics, Kruskal–Wallis analysis of variance was conducted to detect any significant variation among replicates. If no significant variation was detected, the results were pooled for further comparison. Differences between continuous variables were assessed with a non-parametric Mann–Whitney test. The PFU, genomic copies and RNA:PFU ratios were analyzed using non-transformed values. The weight loss data are shown as mean ± standard deviation and statistically analyzed using two-way ANOVA Turkey’s multiple comparison. Analyses were performed in Prism version 9.0 (GraphPad, San Diego, CA).

For virus competition experiments, relative replicative fitness values for mutant virus compared to G614 wt virus were analyzed according to w=(f0/i0), where i0 is the initial mutant:wt ratio and f0 is the final mutant:wt ratio after competition. Sanger sequencing (initial timepoint T0) counts for each virus strain being compared were based upon average counts over three replicate samples of inocula per experiment, and post-infection (timepoint T1) counts were taken from samples of individual subjects. For the primary human airway samples, multiple experiments were performed, so that f0/i0 was clustered by experiment. To model f0/i0, the ratio T0/T1 was found separately for each subject in each strain group, log (base-10) transformed to an improved approximation of normality, and modeled by analysis of variance with relation to group, adjusting by experiment when appropriate to control for clustering within experiment. Specifically, the model was of the form Log10_CountT1overCountT0 ~ Experiment + Group. Fitness ratios between the two groups [the model’s estimate of w=(f0/i0)] were assessed per the coefficient of the model’s Group term, which was transformed to the original scale as 10^coefficient. This modeling approach compensates for any correlation due to clustering within experiment similarly to that of corresponding mixed effect models, and is effective since the number of experiments was small. Statistical analyses were performed using R statistical software (R Core Team, 2019, version 3.6.1). In all statistical tests, two-sided alpha=.05. Catseye plots^29^, which illustrate the normal distribution of the model-adjusted means, were produced using the “catseyes” package^30^.

## Data availability

Extended Data and source data for generating main figures are available in the online version of the paper. Any other information is available upon request.

## Acknowledgments

This research was supported by grants from the NIA and NIAID of the NIH [AI153602 and AG049042 to V.D.M.; R24AI120942 to S.C.W.] and by a STARs Award provided by the University of Texas System to V.D.M.; and Welch Foundation grant AU-0042-20030616 and Cancer Prevention and Research Institute of Texas (CPRIT) Grants RP150551 and RP190561 to Z.A.. P.-Y.S. was supported by NIH grants AI134907, and UL1TR001439, and the authors were also supported by awards from the Sealy & Smith, Kleberg, John S. Dunn, Amon G. Carter, Gilson Longenbaugh and Summerfield Robert Foundations. J.L. was supported by the James W. McLaughlin Fellowship Fund.

## Author contributions

Conceptualization, Y.L., J.L., K.S.P., J.A.P., X.X., V.D.M., P.-Y.S., S.C.W.; Methodology, Y.L., J.L., J.A.P., X.X., X.Z., Z.K., Z.A., D.S., C.S., V.D.M., K.S.P., P.-Y.S., S.C.W.; Investigation, Y.L., J.L., K.S.P., J.A.P., V.D.M., P.-Y.S., S.C.W.; Resources, X.X., Z.K., Z.A.; Data Curation, Y.L., J.L., K.S.P., J.A.P., V.D.M., P.-Y.S., S.C.W.; Writing-Original Draft, P.-Y.S, S.C.W.; Writing-Review & Editing, Y.L., J.L., K.S.P., J.A.P., V.D.M., P.-Y.S., S.C.W.; Supervision, P.-Y.S., S.C.W.; Funding Acquisition, Z.A., V.D.M., P.-Y.S., S.C.W.

---

**Extended Data Figure 1.**
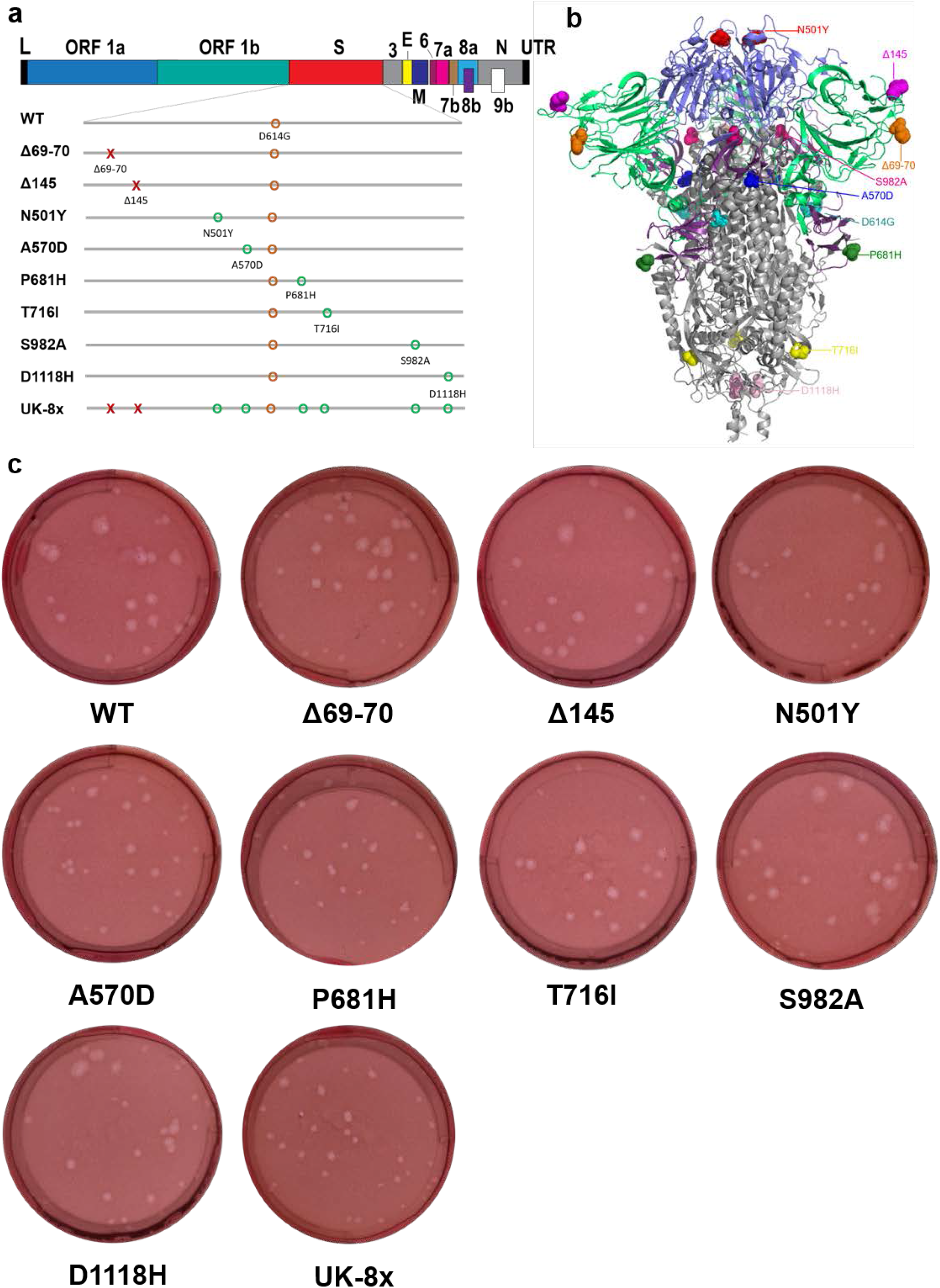
The construction of mutants and their plaque morphologies. **a,** The reverse genetic construction design of all the individual and combined mutations on the wt D614G backbone. Numbers on the upper genome map refer to open reading frames (ORFs). E, envelope glycoprotein gene; L, leader sequence; M, membrane glycoprotein gene; N, nucleocapsid gene; UTR, untranslated region. **b,** The location of all 8 UK B.1.1.7 substitutions and D614G on the SARS-CoV-2 spike protein trimer. **c,** The morphologies of all the rescued mutant SARS-CoV-2 variants. The plaques were stained 2.5 days post-infection of Vero E6 cells.

**Extended Data Figure 2.**
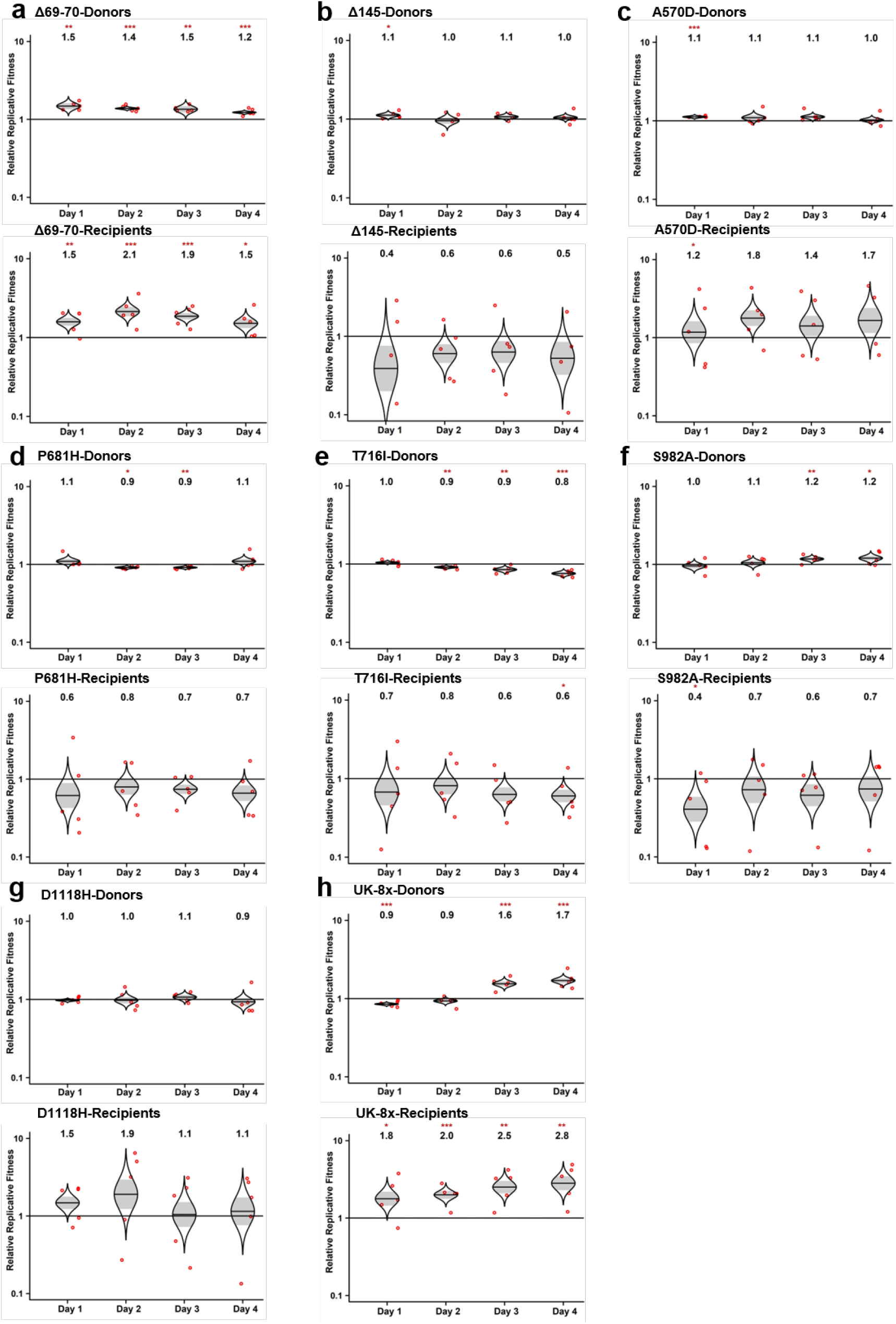
The competition of other SARS-CoV-2 mutants with wt in hamsters. **a-h,** Eight SARS-CoV-2 spike mutants: Δ69-70 (**a**), Δ145 (**b**), A570D (**c**), P681H (**d**), T716I (**e**), S982A (**f**), D1118H (**g**) and UK-8x (**h**) were mixed with wt virus at a ratio of approximately 1:1. The mixture was then inoculated intranasally into donor hamsters and transmitted to the recipient hamsters following the scheme in Figure 1a. The total titer for infection was 10^5^ PFU per hamster. The ratios of the mutant:wt in the nasal washes of hamsters sampled 1-4 days after infection were estimated by Sanger sequencing. Red dots represent individual animals (n = 5), the horizontal lines in each catseye represent the mean, shaded regions represent standard error of the mean; y-axes use a log10 scale. Black numbers above each set of values (catseye) indicate the relative fitness estimates. P values are calculated for the group (strain) coefficient for each linear regression model. *p<0.05; **p<0.01; ***p<0.001.

**Extended Data Figure 3.**
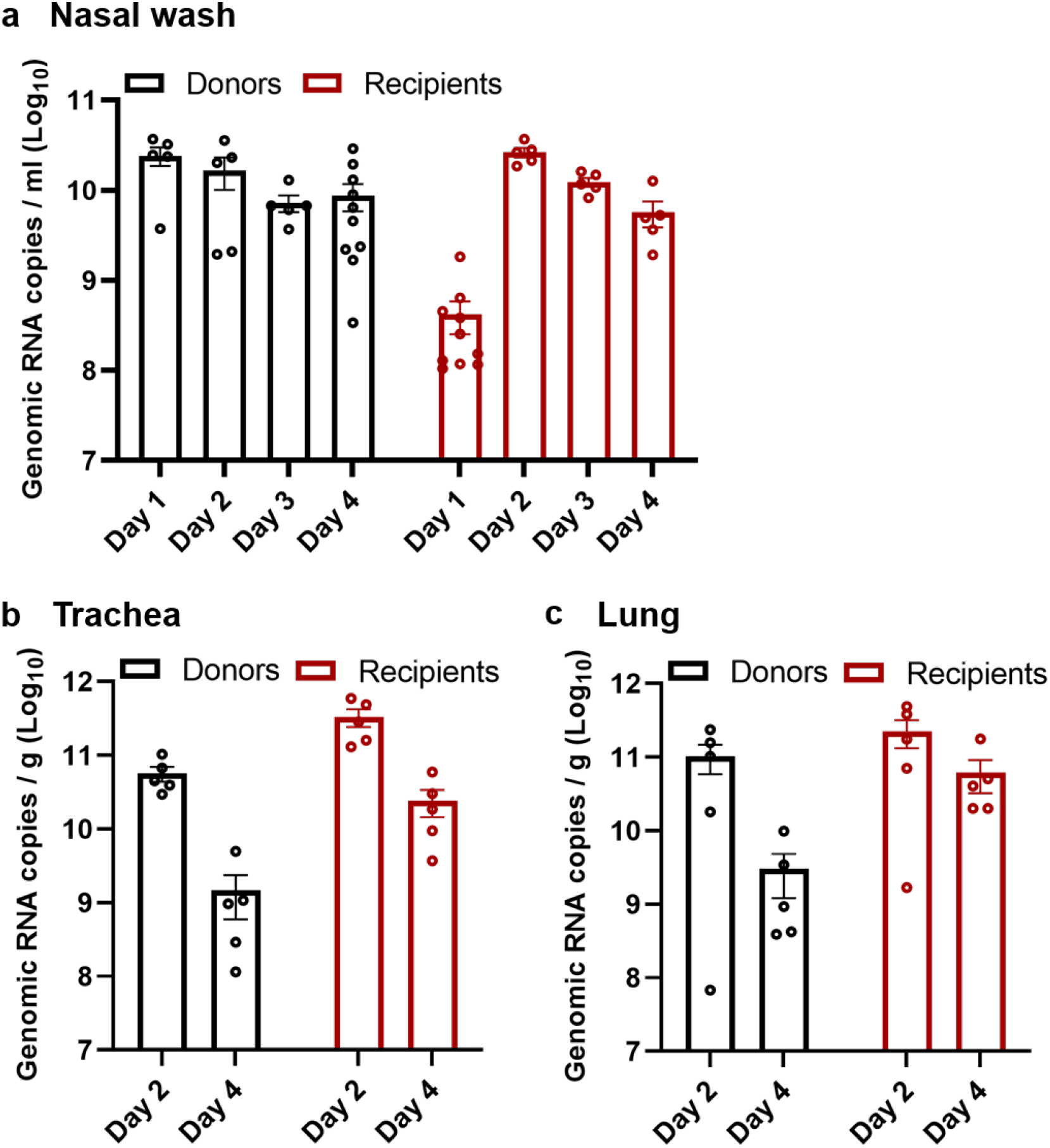
The replication kinetics of N501Y/wt mixed viruses in the competition assay in hamsters. **a-c,** The N501Y variant was mixed with wt virus and inoculated intranasally into hamsters. The replication kinetics of the mixed viruses in the nasal washes (**a**), tracheae (**b**) and lungs (**c**) from both donor and recipient hamsters were measured by real-time qPCR. The nasal wash samples were collected from 1-4 days post-inoculation (donors) or post-contact (recipients). The organ titers in tracheae and lungs from donors and recipients were determined at 2- or 4-days post-inoculation or post-contact. Dots represent individual hamsters (n=10 for nasal wash, n=5 for organ titers). The values in the graph represent the mean ± standard error of the mean.

**Extended Data Figure 4.**
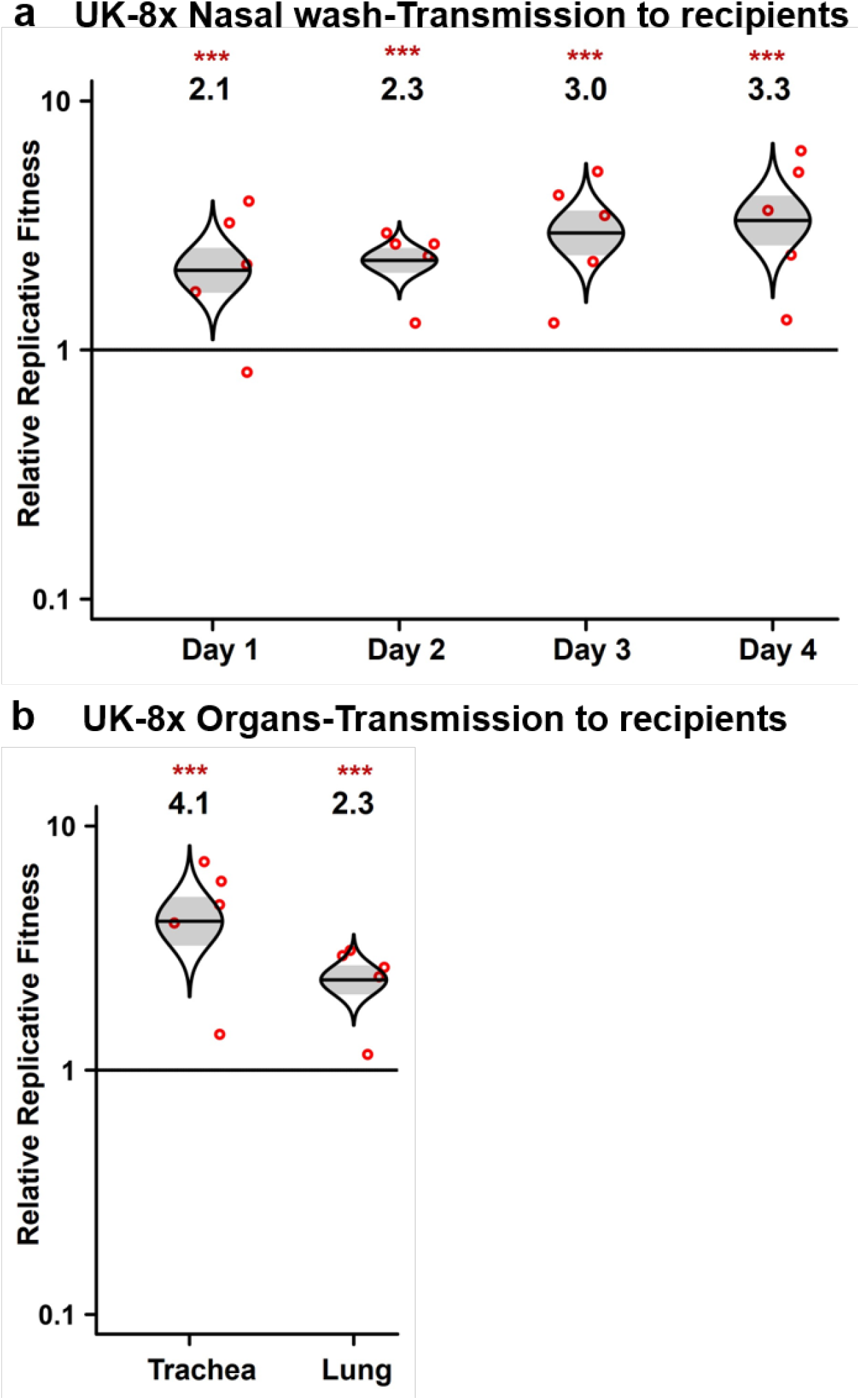
The advantage of the UK-8x mutant during the transmission from donor to recipient hamsters. **a,b,** The ratios of mixed viruses in the nasal washes (**a**), tracheae and lungs (**b**) of recipient hamsters were compared to the ratios of N501Y:wt measured on the day 1 nasal wash of donor hamsters to assess fitness for transmission to and early replication in the recipient hamsters. The total infection titer of the mixed viruses was 10^5^ PFU per hamster. Red dots represent individual animals (n=5), the horizontal lines in each catseye represent the mean, shaded regions represent standard error of the mean; y-axes use a log10 scale. Black numbers above each set of values (catseye) indicate the relative fitness estimates. P values are calculated for the group (strain) coefficient for each linear regression model. ***p<0.001.

**Extended Data Figure 5.**
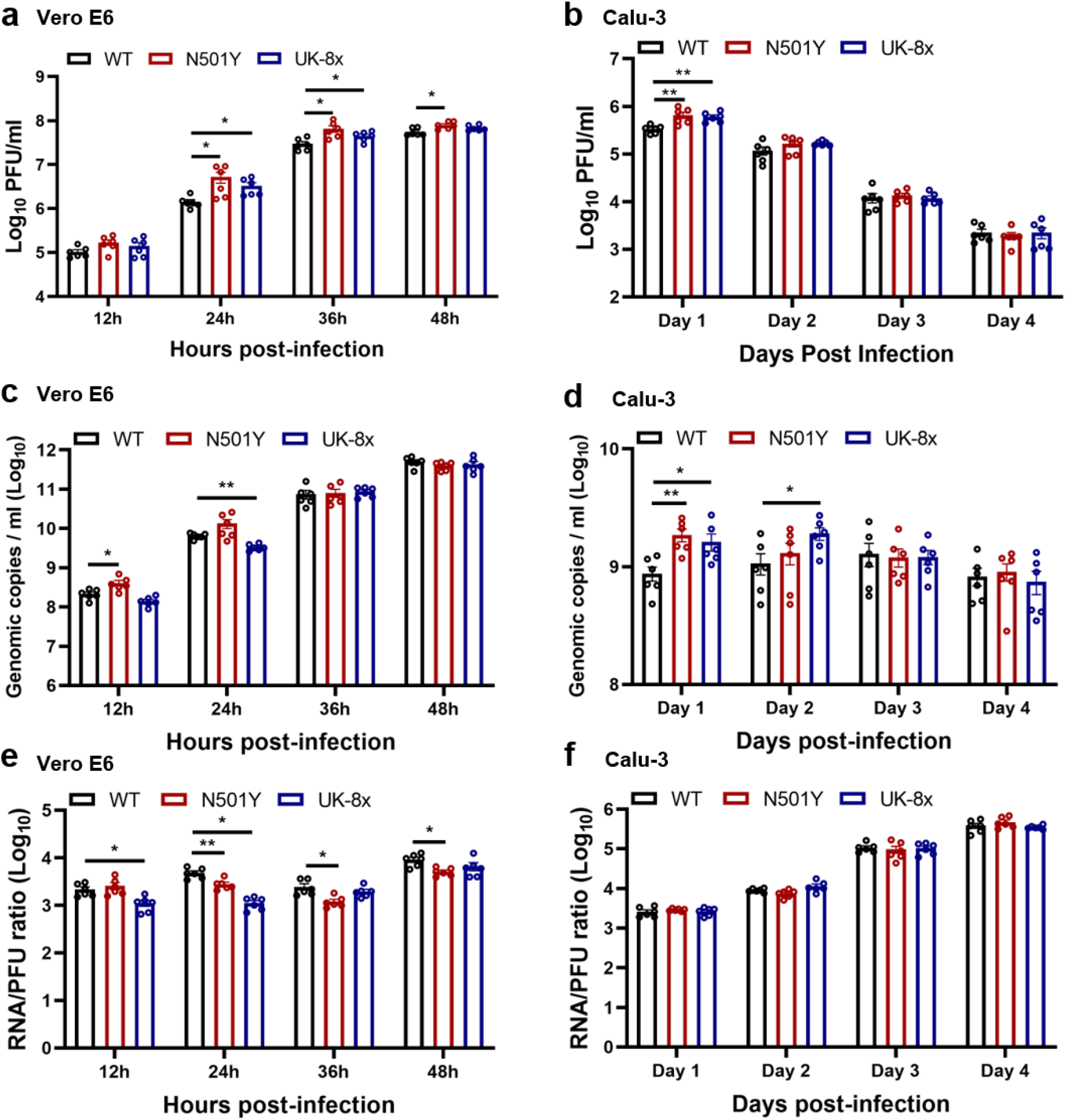
The replication kinetics of the N501Y and UK-8x mutants on Vero E6 and calu-3 cells. **a-f,** The wt, N501Y and UK-8x mutant viruses were inoculated on Vero E6 (**a,c,e**) and calu-3 cells (**b,d,f**) respectively. The amounts of infectious virus (**a,b**) and genomic RNA (**c,d**) were quantified by plaque assay and RT–qPCR, respectively. The genomic RNA:PFU ratio (**e,f**) was calculated as an indication of virion infectivity. The Vero E6 and calu-3 were infected at a MOI of 0.01 and 0.1, respectively. The detection limit of the plaque assay was 10 PFU/ml. Dots represent individual biological replicates (n=6) pooled from two independent experiments. The values in the graph represent the mean ± standard error of the mean. A non-parametric Mann-Whitney test was used to determine significant differences. P values were adjusted using the Bonferroni correction to account for multiple comparisons. Differences were considered significant if *p<0.025; **p<0.005.

**Extended Data Figure 6.**
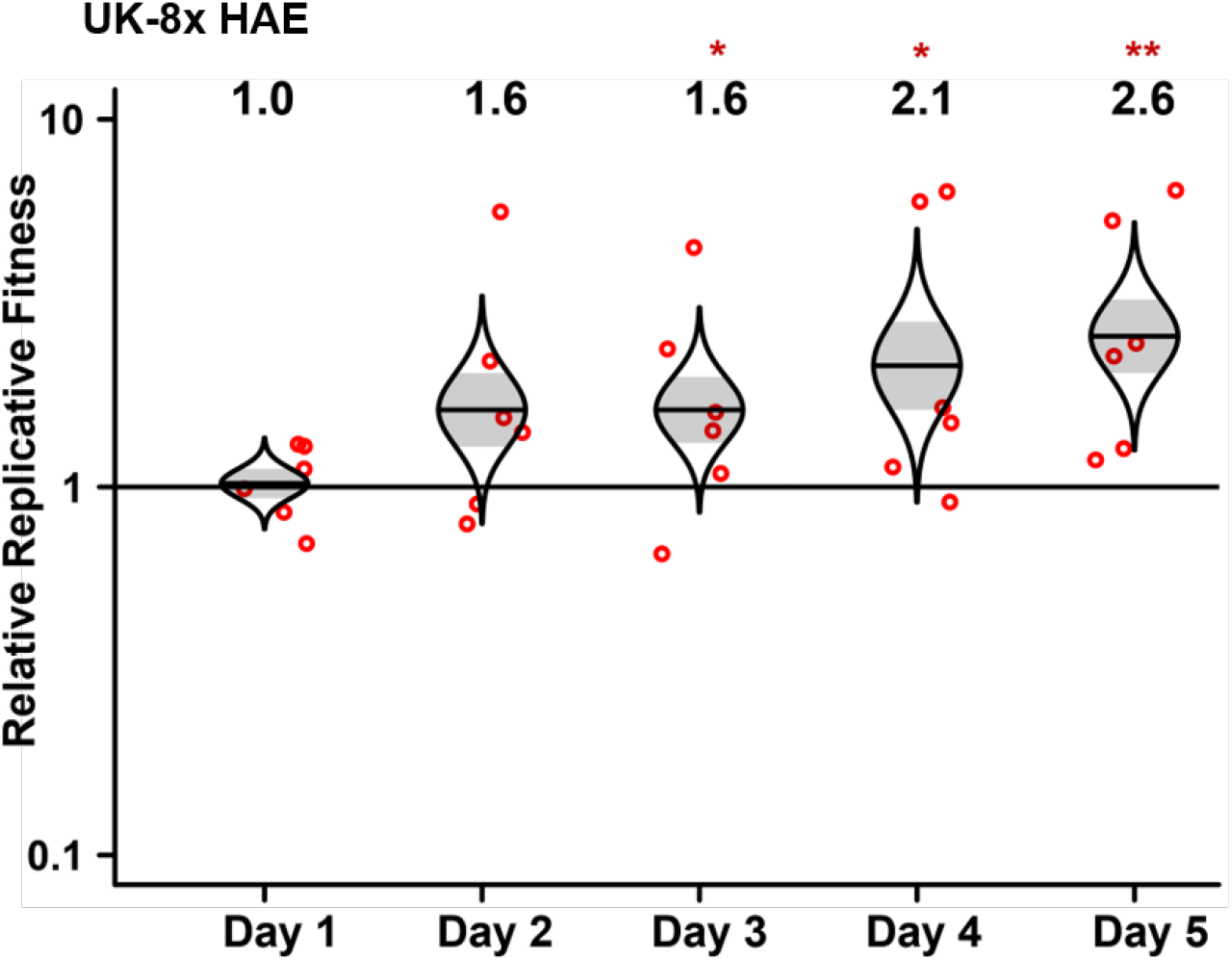
The competition assay of UK-8x against wt on primary human airway epithelial cells. The UK-8x variant was mixed with wt virus and inoculated on the human airway epithelial (HAE) cells at a total MOI of 5. The ratios of UK-8x mutant to the wt virus were measured by Sanger sequencing. Red dots represent individual biological replicates (n=6), pooled from 2 independent experiments. The horizontal lines in each catseye represent the mean, shaded regions represent standard error of the mean; y-axes use a log10 scale. Black numbers above each set of values (catseye) indicate the relative fitness estimates. P values are calculated for the group (strain) coefficient for each linear regression model. *p<0.05; **p<0.01.

**Extended Data Figure 7.**
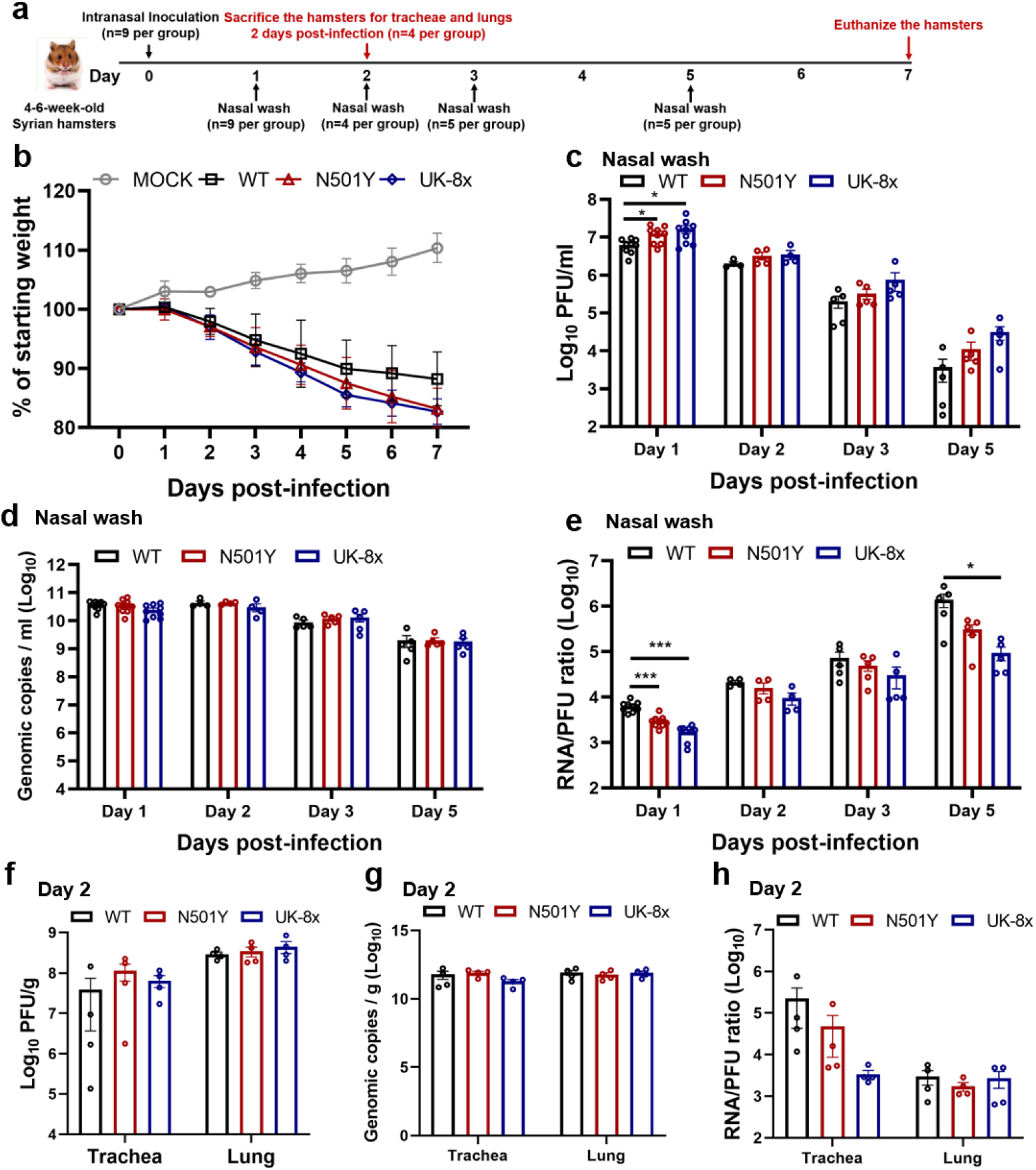
The spike N501Y substitution benefits viral infection of hamster upper airways. **a,** Design of the hamster infection kinetic studies. The wt, N501Y and UK-8x viruses were intranasally inoculated into hamsters at a titer of 10^4^ PFU per hamster. Nine hamsters were utilized for the initial infection in each group. At 2 days-post infection, 4 hamsters were sacrificed for organ collections. The nasal washes of the hamsters were collected on days 1, 3 and 5 post-infection or before sacrifice. **b,** Weight change in hamsters following infection by the N501Y (n=5) and UK-8x (n=5) mutants compared to the wt (n=5). MOCK group (n=4) served as the negative (uninfected) control. The body weights were measured form 1-7 days post-infection. The weight loss data are shown as mean ± standard deviation and statistically analyzed using two-way ANOVA Turkey’s multiple comparison. No significant differences were seen between the N501Y/UK-8x and wt groups. **c-h,** The infection of N501Y and UK-8x mutants compared to the wt in the nasal washes (**c-e**) collected 1, 2, 3, or 5 days post-infection and in the organs (**f-h**) 2 days post-infection. The amounts of infectious virus (**c,f**) and genomic RNA (**d,g**) were quantified by plaque assay and RT–qPCR, respectively. The genomic RNA:PFU ratio (**e,h**) was calculated as an indication of virion infectivity. The detection limitation of the plaque assay was 10 PFU/ml. The values in the graph represent the mean ± standard error of the mean. A non-parametric Mann-Whitney test was used to determine significant differences. P values were adjusted using the Bonferroni correction to account for multiple comparisons. Differences were considered significant if *p<0.025; ***p<0.0005.

**Extended Data Table 1.**
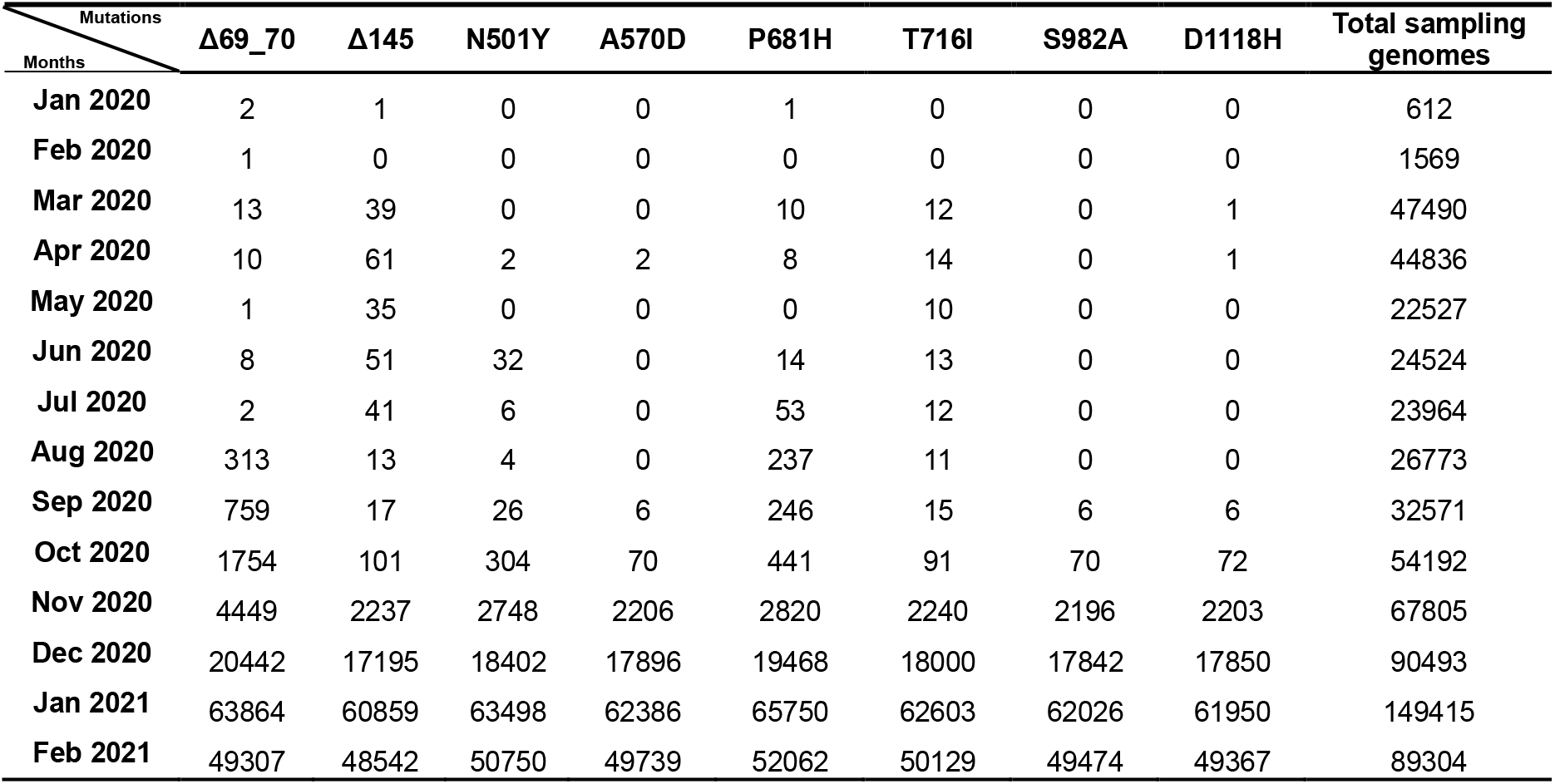
Total numbers of the 8 UK spike substitutions in sequenced SARS-CoV-2 genomes.

**Extended Data Table 2.**
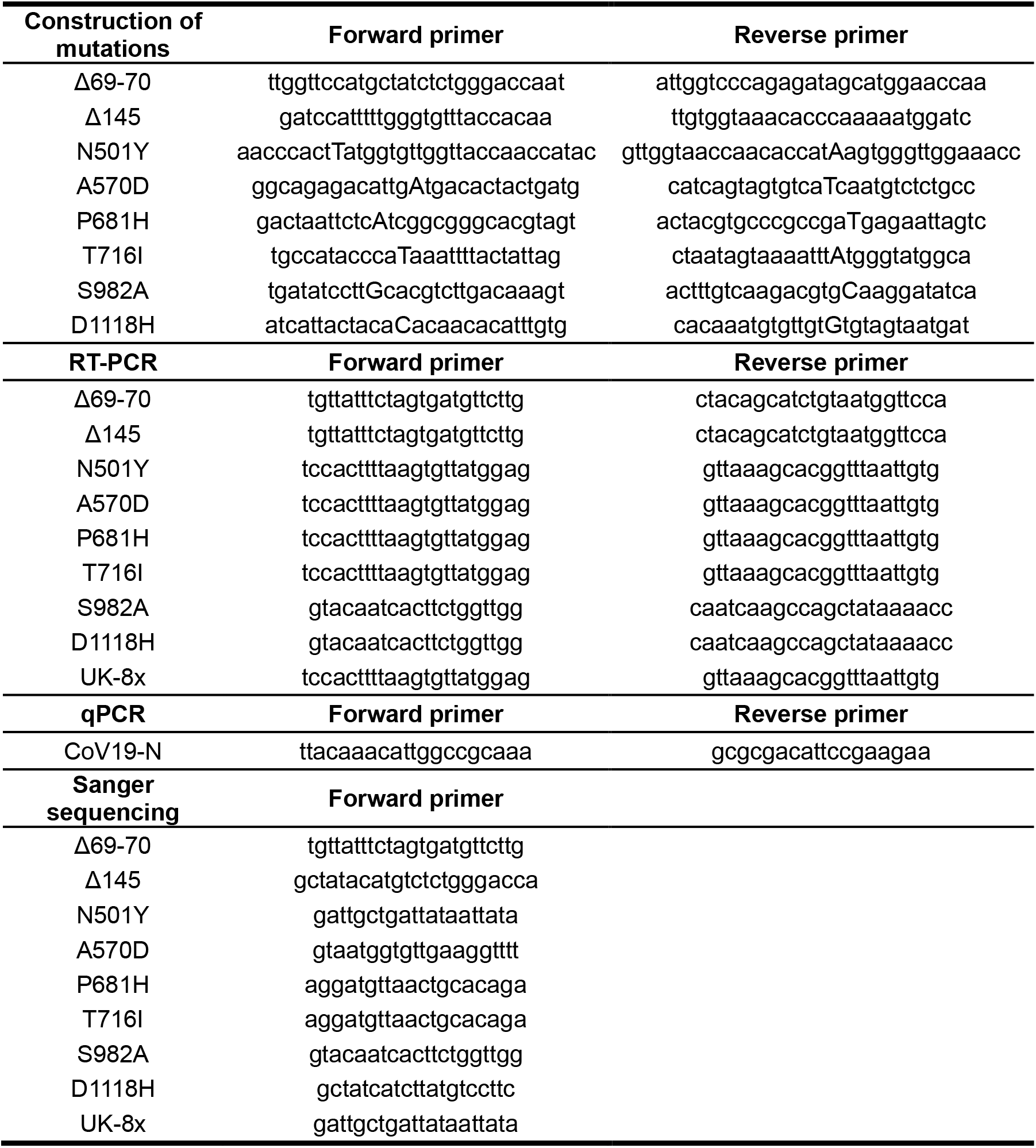
Primer list used for mutation construction, RT-PCR, qPCR and sanger sequencing.

## References

1 Zhou, P. et al. A pneumonia outbreak associated with a new coronavirus of probable bat origin. Nature 579, 270–273, doi:10.1038/s41586-020-2012-7 (2020).

2 Smith, E. C., Blanc, H., Surdel, M. C., Vignuzzi, M. & Denison, M. R. Coronaviruses lacking exoribonuclease activity are susceptible to lethal mutagenesis: evidence for proofreading and potential therapeutics. PLoS Pathog 9, e1003565, doi:10.1371/journal.ppat.1003565 (2013).

3 Gribble, J. et al. The coronavirus proofreading exoribonuclease mediates extensive viral recombination. PLoS Pathog 17, e1009226, doi:10.1371/journal.ppat.1009226 (2021).

4 Huang, Y., Yang, C., Xu, X. F., Xu, W. & Liu, S. W. Structural and functional properties of SARS-CoV-2 spike protein: potential antivirus drug development for COVID-19. Acta pharmacologica Sinica 41, 1141–1149, doi:10.1038/s41401-020-0485-4 (2020).

5 Starr, T. N. et al. Deep Mutational Scanning of SARS-CoV-2 Receptor Binding Domain Reveals Constraints on Folding and ACE2 Binding. Cell 182, 1295–1310 e1220, doi:10.1016/j.cell.2020.08.012 (2020).

6 Harcourt, J. et al. Severe Acute Respiratory Syndrome Coronavirus 2 from Patient with 2019 Novel Coronavirus Disease, United States. Emerg Infect Dis 26, doi:10.3201/eid2606.200516 (2020).

7 Hou, Y. J. et al. SARS-CoV-2 D614G variant exhibits efficient replication ex vivo and transmission in vivo. Science 370, 1464–1468, doi:10.1126/science.abe8499 (2020).

8 Plante, J. A. et al. Spike mutation D614G alters SARS-CoV-2 fitness. Nature, doi:10.1038/s41586-020-2895-3 (2020).

9 Chan, J. F. et al. Simulation of the Clinical and Pathological Manifestations of Coronavirus Disease 2019 (COVID-19) in a Golden Syrian Hamster Model: Implications for Disease Pathogenesis and Transmissibility. Clin. Infect. Dis. 71, 2428–2446, doi:10.1093/cid/ciaa325 (2020).

10 Zhou, B. et al. SARS-CoV-2 spike D614G change enhances replication and transmission. Nature, doi:10.1038/s41586-021-03361-1 (2021).

11 Leung, K., Shum, M. H., Leung, G. M., Lam, T. T. & Wu, J. T. Early transmissibility assessment of the N501Y mutant strains of SARS-CoV-2 in the United Kingdom, October to November 2020. Euro Surveill 26, doi:10.2807/1560-7917.ES.2020.26.1.2002106 (2021).

12 Galloway, S. E. et al. Emergence of SARS-CoV-2 B.1.1.7 Lineage - United States, December 29, 2020-January 12, 2021. MMWR Morb Mortal Wkly Rep 70, 95–99, doi:10.15585/mmwr.mm7003e2 (2021).

13 Claro, I. M. et al. Local Transmission of SARS-CoV-2 Lineage B.1.1.7, Brazil, December 2020. Emerg Infect Dis 27, doi:10.3201/eid2703.210038 (2021).

14 Chen, R. E. et al. Resistance of SARS-CoV-2 variants to neutralization by monoclonal and serum-derived polyclonal antibodies. Nat. Med., doi:10.1038/s41591-021-01294-w (2021).

15 Xie, X. et al. Neutralization of SARS-CoV-2 spike 69/70 deletion, E484K and N501Y variants by BNT162b2 vaccine-elicited sera. Nat. Med., doi:10.1038/s41591-021-01270-4 (2021).

16 Liu, Y. et al. Neutralizing Activity of BNT162b2-Elicited Serum - Preliminary Report. N Engl J Med, doi:10.1056/NEJMc2102017 (2021).

17 Harcourt, J. et al. Severe Acute Respiratory Syndrome Coronavirus 2 from Patient with Coronavirus Disease, United States. Emerg Infect Dis 26, 1266–1273, doi:10.3201/eid2606.200516 (2020).

18 Xie, X. et al. An Infectious cDNA Clone of SARS-CoV-2. Cell Host Microbe, doi:10.1016/j.chom.2020.04.004 (2020).

19 Wiser, M. J. & Lenski, R. E. A Comparison of Methods to Measure Fitness in Escherichia coli. PLoS One 10, e0126210, doi:10.1371/journal.pone.0126210 (2015).

20 Grubaugh, N. D. et al. Genetic Drift during Systemic Arbovirus Infection of Mosquito Vectors Leads to Decreased Relative Fitness during Host Switching. Cell Host Microbe 19, 481–492, doi:10.1016/j.chom.2016.03.002 (2016).

21 Bergren, N. A. et al. “Submergence” of Western equine encephalitis virus: Evidence of positive selection argues against genetic drift and fitness reductions. PLoS Pathog 16, e1008102, doi:10.1371/journal.ppat.1008102 (2020).

22 Coffey, L. L. & Vignuzzi, M. Host alternation of chikungunya virus increases fitness while restricting population diversity and adaptability to novel selective pressures. J. Virol. 85, 1025–1035, doi:10.1128/JVI.01918-10 (2011).

23 Liu, J. et al. Role of mutational reversions and fitness restoration in Zika virus spread to the Americas. Nat Commun 12, 595, doi:10.1038/s41467-020-20747-3 (2021).

24 Kemp, S. A. et al. Recurrent emergence and transmission of a SARS-CoV-2 Spike deletion ΔH69/ΔV70. bioRxiv (2020).

25 Chan, J. F. et al. Simulation of the clinical and pathological manifestations of Coronavirus Disease 2019 (COVID-19) in golden Syrian hamster model: implications for disease pathogenesis and transmissibility. Clin. Infect. Dis., doi:10.1093/cid/ciaa325 (2020).

26 Xie, X. et al. Engineering SARS-CoV-2 using a reverse genetic system. Nat Protoc, doi:10.1038/s41596-021-00491-8 (2021).

27 Carr, I. M. et al. Inferring relative proportions of DNA variants from sequencing electropherograms. Bioinformatics 25, 3244–3250, doi:10.1093/bioinformatics/btp583 (2009).

28 Ku, Z. et al. Molecular determinants and mechanism for antibody cocktail preventing SARS-CoV-2 escape. Nat Commun 12, 469, doi:10.1038/s41467-020-20789-7 (2021).

29 Cumming, G. The New Statistics: Why and How. Psychol. Sci., 7–29, doi:10.1177/0956797613504966 (2014).

30 Andersen, C. Catseyes: Create Catseye Plots Illustrating the Normal Distribution of the Means. R package version 0.2.3. (2019).

